# Replication kinetic, cell tropism and associated immune responses in SARS-CoV-2 and H5N1 virus infected human iPSC derived neural models

**DOI:** 10.1101/2021.03.15.435472

**Authors:** Lisa Bauer, Bas Lendemeijer, Lonneke Leijten, Carmen W. E. Embregts, Barry Rockx, Steven A. Kushner, Femke M.S. de Vrij, Debby van Riel

**Affiliations:** Department of Viroscience, Erasmus Medical Center, Rotterdam, The Netherlands; Department of Psychiatry, Erasmus Medical Center, Rotterdam, The Netherlands

**Keywords:** neurotropism, hiPSC neurons, coronavirus, SARS-CoV-2, COVID-19, influenza A virus, H5N1 virus, IL-8, interferon

## Abstract

Severe acute respiratory syndrome coronavirus-2 (SARS-CoV-2) infection is associated with a wide variety of neurological complications. Even though SARS-CoV-2 is rarely detected in the central nervous system (CNS) or cerebrospinal fluid, evidence is accumulating that SARS-CoV-2 might enter the CNS via the olfactory nerve. However, what happens after SARS-CoV-2 enters the CNS is poorly understood. Therefore, we investigated the replication kinetics, cell tropism, and associated immune responses of SARS-CoV-2 infection in different types of neural cultures derived from human induced pluripotent stem cells (hiPSCs). SARS-CoV-2 was compared to the neurotropic and highly pathogenic H5N1 influenza A virus. SARS-CoV-2 infected a minority of individual mature neurons, without subsequent virus replication and spread, despite ACE2, TMPRSS2 and NPR1 expression in all cultures. However, this sparse infection did result in the production of type-III-interferons and IL-8. In contrast, H5N1 virus replicated and spread very efficiently in all cell types in all cultures. Taken together, our findings support the hypothesis that neurological complications might result from local immune responses triggered by virus invasion, rather than abundant SARS-CoV-2 replication in the CNS.

## Introduction

Neurological manifestations are present in a substantial proportion of patients suffering from the respiratory coronavirus disease 2019 (COVID-19). Symptoms comprise loss of smell (anosmia), loss of taste (hypogeusia), headache, fatigue, nausea and vomiting^1–3^. Additionally, more severe neurological complications such as seizures, confusion, cerebrovascular injury, stroke, encephalitis, encephalopathies and altered mental status are being increasingly reported in hospitalized patients^4–6^.

It remains to be established whether the reported neurological manifestations are a direct consequence of local invasion of severe acute respiratory syndrome coronavirus-2 (SARS-CoV-2) into the central nervous system (CNS), an indirect consequence of the associated systemic immune responses, or a combination of both. In human and animal models, it has been shown that SARS-CoV-2 is able to replicate in the olfactory mucosa^7,8^, suggesting that the olfactory nerve could function as an important route of entry into the CNS^9^, as observed previously for other respiratory viruses^10^. Post-mortem brain tissue analyses of fatal COVID-19 cases has revealed mild neuropathological changes which might be related to hypoxia-and pronounced neuroinflammation in different regions of the brain^11^. In the majority of cases, neither SARS-CoV-2 viral RNA, nor virus antigen, could be detected in the CNS^5,12^. In line with this, SARS-CoV-2 viral RNA has rarely been detected in the cerebrospinal fluid (CSF) of COVID-19-patients with neurological symptoms^13–15^. Together, this suggests that SARS-CoV-2 might enter the CNS but unable to replicate there efficiently.

In the brain, viruses encounter a variety of different cell types such as neurons, astrocytes and microglia. Investigations of CNS cell-type specific infection of SARS-CoV-2 have been inconsistent^16–23^. Most of studies have investigated virus replication by detection of viral RNA, but have not reported whether infectious progeny viruses are produced. Therefore, in order to investigate replication and infection efficiency, cell tropism and associated immune responses of SARS-CoV-2, we differentiated human induced pluripotent stem cells (hiPSCs) along a variety of different neural lineage specifications, which afforded a unique and flexible platform to study the neurotropism of viruses *in vitro*. Specifically, we directed hiPSC-colonies towards embryoid bodies with differentiation into neural progenitor cells (NPCs) and subsequently mature neural networks^24^. In addition, we also utilized a rapid neuronal differentiation protocol based on forced overexpression of the transcription factor Ngn2 in hiPSCs (Figure 1A) ^25,26^ to generate pure populations of neurons that we co-cultured with hiPSC-derived astrocytes. Using these specified CNS cell types, we directly compared the characteristics of SARS-CoV-2 with the highly pathogenic H5N1 influenza A virus, a virus with zoonotic potential which is known to efficiently replicate in neural cells *in vivo*^27–32^ and *in vitro*^33–36^.

**Figure 1.**
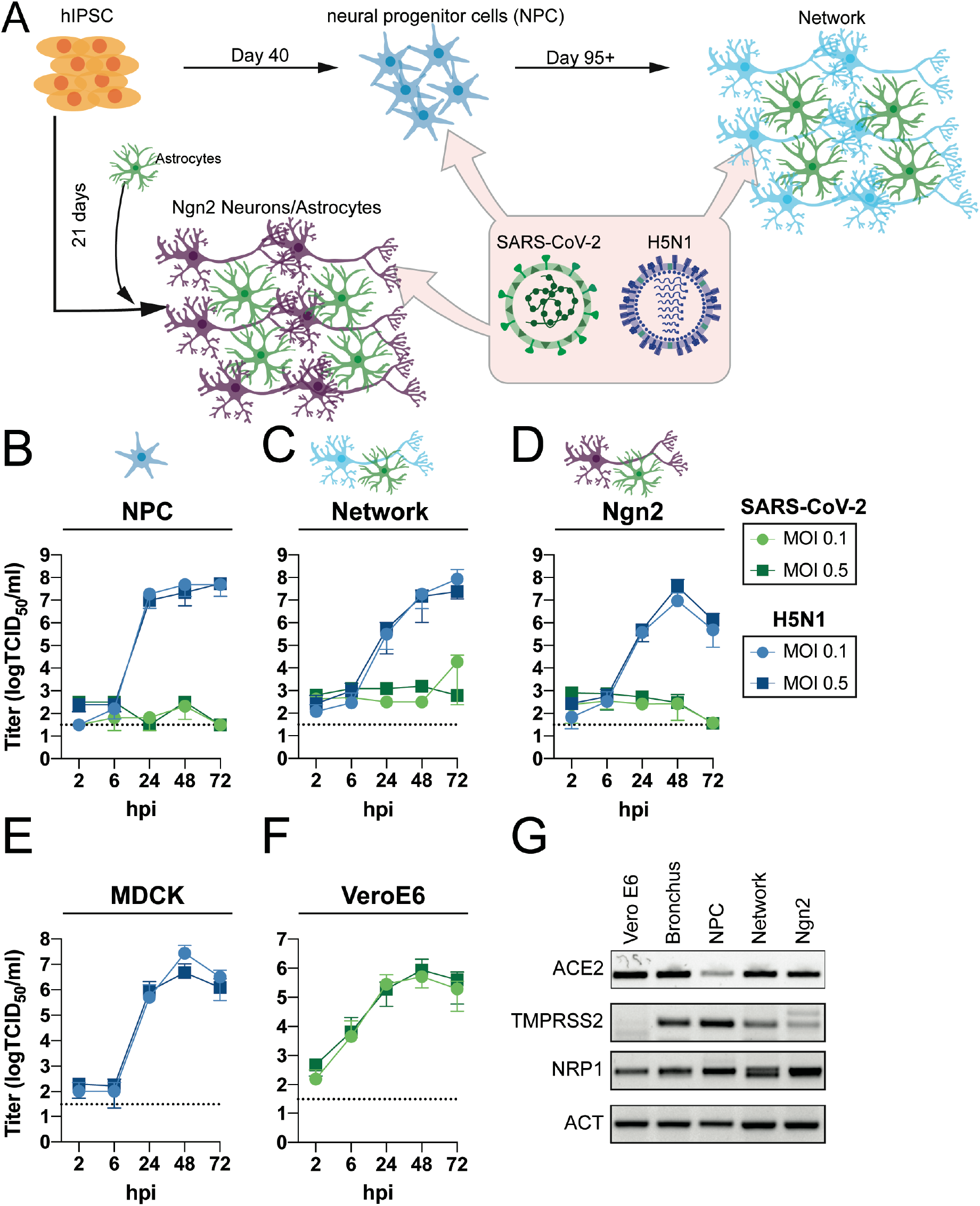
SARS-CoV-2 does not replicate in hiPSC derived neural (co-)cultures in contrast to H5N1 virus. (A) A schematic depiction of the different hiPSC-derived differentiation strategies of the neural cultures. hiPSC are differentiated into neural progenitor cells (NPC) and subsequently into mixed neural cultures containing mixed neurons and astrocytes. Alternatively, hiPSC are differentiated into excitatory neurons by inducing overexpression of Ngn2, these are grown in a co-culture with hiPSC-derived astrocytes. Growth kinetics of SARS-CoV-2 or H5N1 virus, in hiPSC-derived (B) NPCs, (C) mature neural networks, or (D) Ngn2 co-cultures using a MOI of 0.1 and 0.5. As positive controls (E) MDCK and (F) VeroE6 cells were infected with H5N1 virus or SARS-CoV2, respectively. Data represent mean ± standard deviation (SD) from three independent experiments. Every growth curve was performed either in biological duplicates or triplicates. (G) Presence of the host factors angiotensin-converting enzyme 2 (ACE2), Transmembrane protease serine 2 (TMPRSS2) and neuropilin-1 (NRP1) of the neural cultures was determined with PCR. As controls for the expression of ACE2 and TMPRSS2 bronchus bronchiole organoids were used. The uncropped agarose gels are displayed in Supplement figure 1.

## Results

### SARS-CoV-2 does not replicate efficiently in hiPSC neural cell types, despite the presence of ACE2, TMPRSS2 and NRP1

To investigate the replication efficiency of SARS-CoV-2, we utilized hiPSC-derived NPCs and differentiated these to mature neural cultures (Figure 1A). NPCs and fully differentiated neural cultures were infected with SARS-CoV-2 and H5N1 virus at a multiplicity of infection (MOI) of 0.1 and 0.5. At 2, 6, 24 48 and 72 hours post infection (hpi), infectious virus titers in the supernatants were determined by endpoint titration. In contrast to H5N1 virus, no productive infection in SARS-CoV-2 inoculated NPC and mature neural cultures was detected (Figure 1B and Figure 1C). As an alternative to the laborious and time-consuming differentiation of mature neural networks through embryoid bodies and NPC stages, we also employed a rapid differentiation protocol that yields a pure culture of iPSC-derived glutamatergic cortical neurons by overexpressing the transcription factor neurogenin-2 (Ngn2)^25^. We further supplemented the Ngn2-induced neurons with hiPSC-derived astrocytes to support their survival and maturation (Figure 1A). SARS-CoV-2 did not replicate efficiently in the Ngn2 co-cultures, in contrast to H5N1 virus (Figure 1D). As a positive control for virus replication, Vero-E6 and MDCK cells were infected with SARS-CoV-2 and H5N1 virus, respectively (Figure 1E and 1F).

Next, we evaluated the presence of important SARS-CoV-2 entry factors, such as angiotensin-converting enzyme 2 (ACE2), transmembrane protease serine 2 (TMPRSS2) and neuropilin-1 (NRP1). In all cultures, there was clear evidence for ACE2, TMPRSS2 and NRP1 expression, suggesting cellular susceptibility to SARS-CoV-2 virus infection (Figure 1G, Supplement Figure 1).

### SARS-CoV-2 infects MAP2-expressing neurons and does not induce caspase-3 expression

To determine whether SARS-CoV-2 was able to infect individual cells, we stained for virus antigen 72 hpi after infection with a MOI of 0.5. SARS-CoV-2 sparsely infected cells in neural cultures at 72 hpi. Infection was observed in single scattered MAP2-positive neurons (Figure 2A and 2B). In one experiment, we identified a cluster of MAP2-positive cells that stained positively for SARS-CoV-2 nuclear protein (NP) at 72 hpi (Supplement Figure 2A). In the Ngn2 co-cultures, we were only able to detect MAP2/NEUN^+^ neurons positive for SARS-CoV-2 NP, suggesting that SARS-CoV-2 infects only mature neurons and does so only sparsely (Supplement Figure 2B). We found no convincing evidence of SARS-CoV-2 NP-positive cells among SOX2^+^ NPCs or GFAP^+^ astrocytes (Figure 2A-2C). H5N1 virus abundantly infected SOX2^+^ NPCs, GFAP^+^ astrocytes and MAP2^+^ neurons (Figure 2A-C).

**Figure 2.**
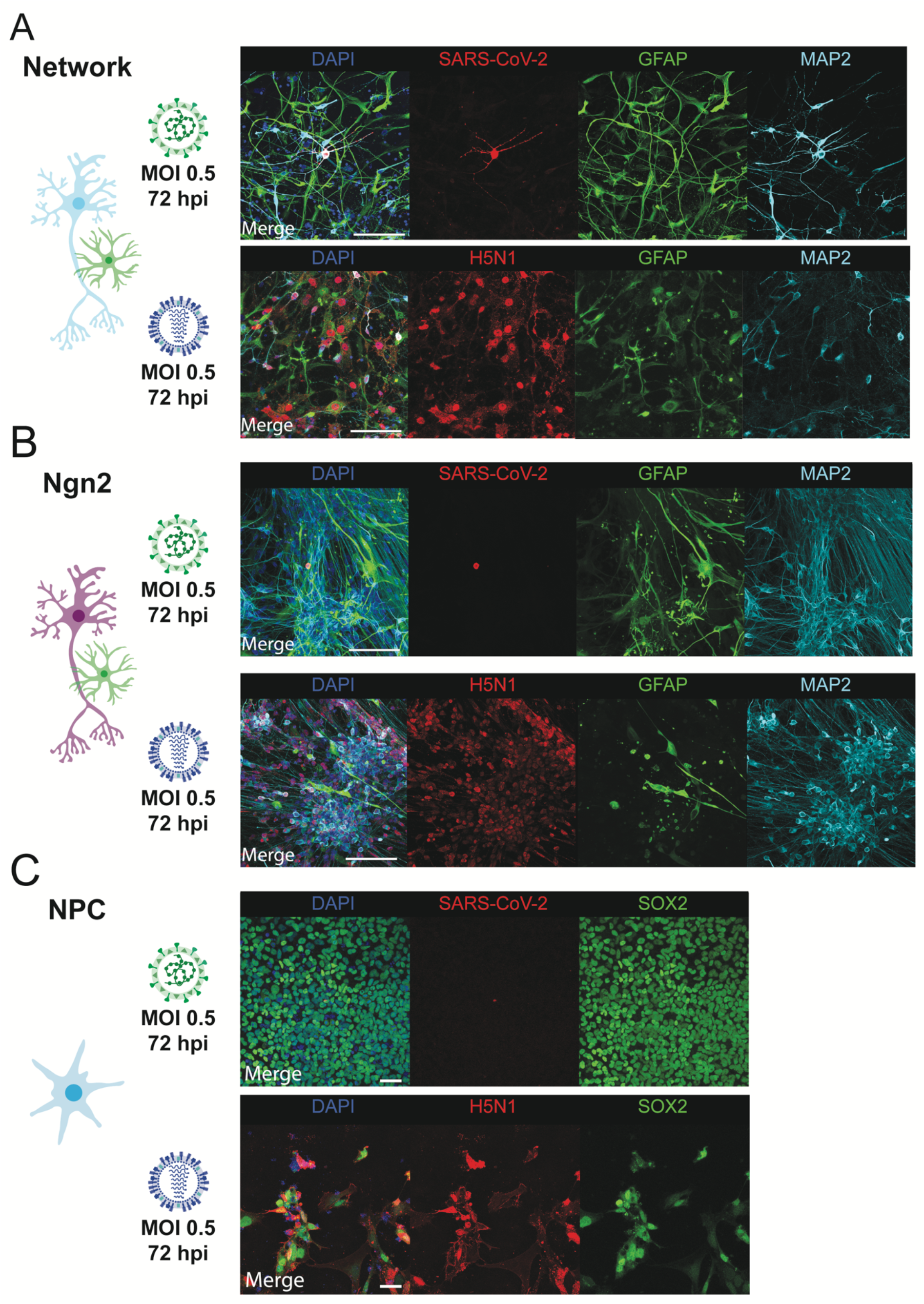
SARS-CoV-2 infects MAP2^+^ neurons. (A) Mixed neural culture (scale bar = 100 μm), (B) Ngn2 co-cultures (scale bar = 100 μm) and (C) NPCs (scale bar = 50 μm) were infected with a MOI of 0.5 with SARS-CoV-2 or H5N1 virus, respectively. 72 hours post infection, the cells were fixed, stained for the presence of viral antigen (SARS-CoV-2 NP or H5N1 NP in red). MAP2 (cyan) was used as a marker for neurons, astrocytes were identified by staining for glial fibrillary acidic protein (GFAP) (green) and SOX2 (green) was used as a marker for NPCs. Cells were counterstained with DAPI (blue) to visualize the nuclei. Data shown are representative examples from three independent experiments for each culture condition.

Next, we wanted to investigate whether SARS-CoV-2 infection induced neuronal apoptosis. Therefore, we infected Ngn2 co-cultures with SARS-CoV-2 and stained for the apoptosis marker caspase-3. We again observed that SARS-CoV-2 infected only MAP2^+^ neurons. Neurons expressing SARS-CoV-2 NP did not exhibit caspase-3 expression (Figure 3A and 3B and Supplement Figure 2C).

**Figure 3.**
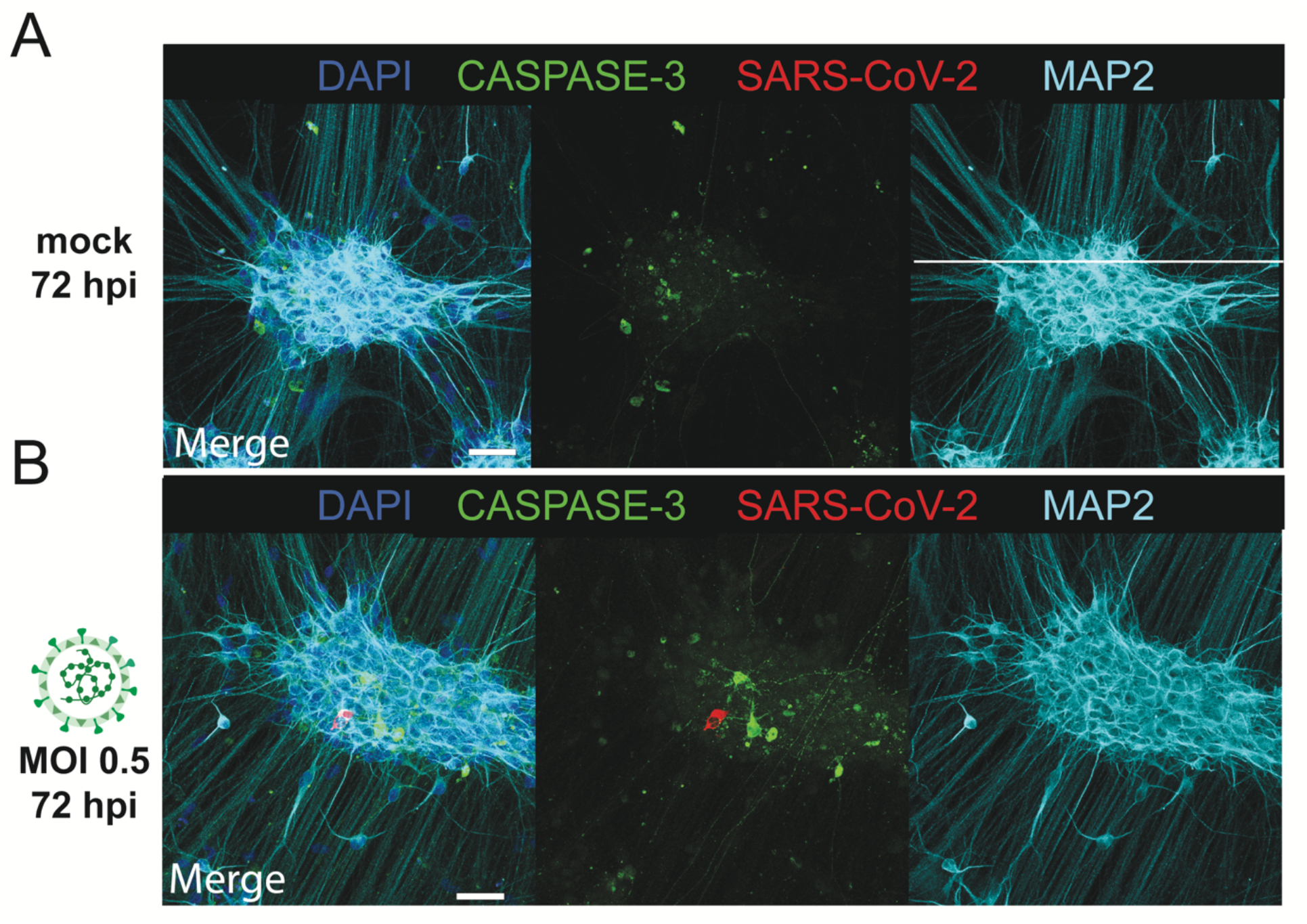
SARS-CoV-2 infections does not result in upregulation of caspase-3. Ngn2 co-cultures were either (A) mock infected or infected with (B) SARS-CoV-2 at a MOI 0.5 (scale bar = 50 μm). 72 hours post infection, the cells were fixed and stained for the presence of SARS-CoV-2 antigen (red) or for the apoptosis marker caspase-3 (green). Data shown are representative examples from two independent experiments.

**Figure 4.**
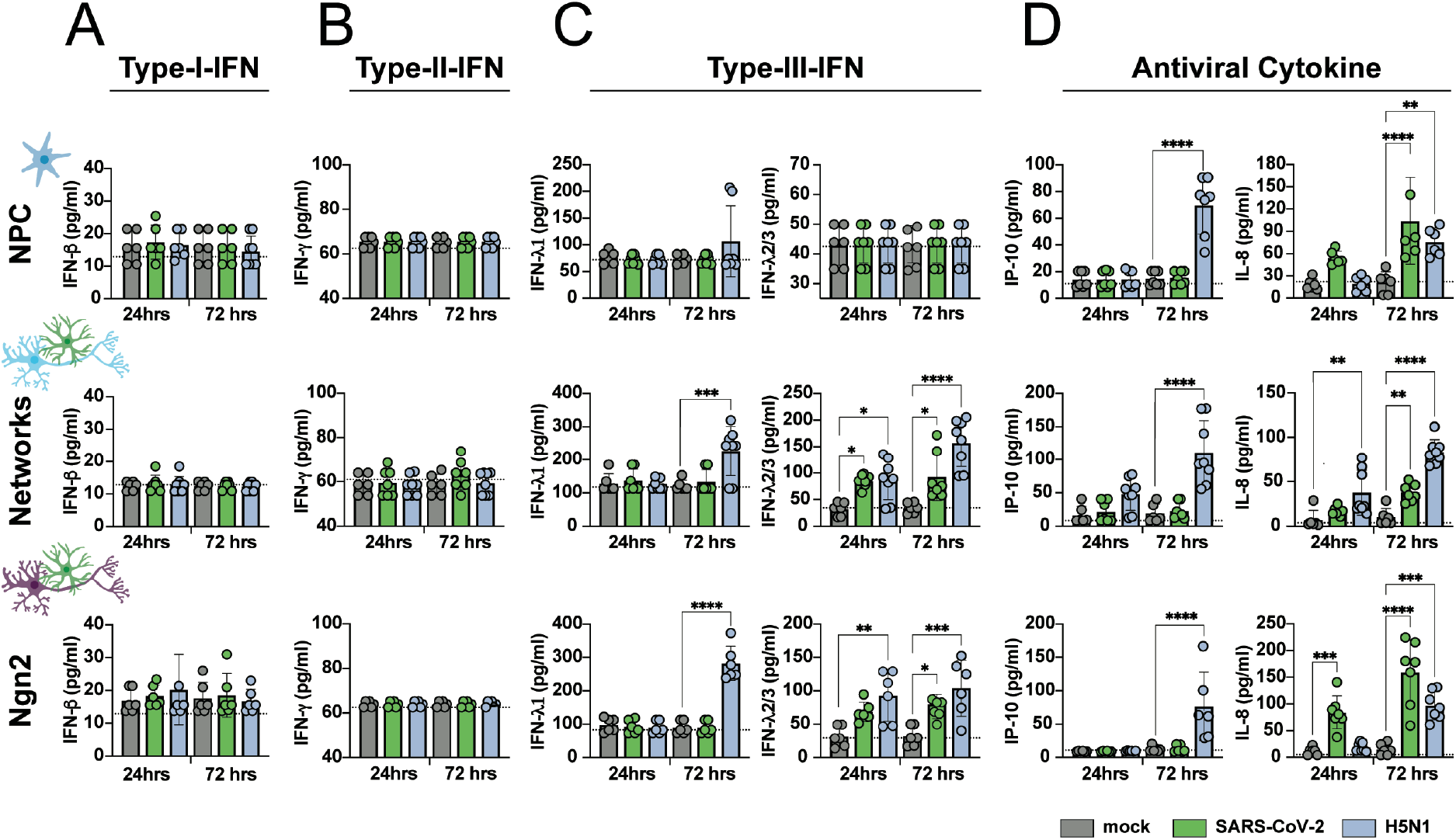
SARS-CoV-2 infection induces type-III-IFN and IL-8. NPCs, mixed neural cultures and Ngn2 co-cultures were infected with SARS-CoV-2 and H5N1 virus at a MOI 0.1. Concentration of (A) type-I-interferon (IFN-β), (B) type-II-IFN (IFN-γ), (C) type-III-IFN (IFNλ1 and IFNλ2/3) and (D) the antiviral cytokines IP-10 and IL-8 were measured in the supernatant 24 and 72 hours post infection. The data are derived from three independent experiments and each experiment was performed either in biological duplicates or triplicates. The assay was performed in technical duplicates for each sample. The data displayed represent average values of the technical duplicates of each experiment performed. Error bars denote mean ± standard deviation (SD). Statistical significance was calculated with a one-way ANOVA with a Bonferroni post hoc test, the means of the mock infected samples were compared to the means of the SARS-CoV-2 and H5N1 virus infected samples at 24 and 72 hours post infection. Asterisks (*) indicate statistical significance. *P < 0.05, **P < 0.01, ***P < 0.001,****P < 0.0001.

### SARS-CoV-2 infection induces IFNλ2/3 and IL-8

To determine the immune response of the neural cultures towards SARS-CoV-2 and H5N1 virus infection, we measured a panel of antiviral cytokines in the supernatant of infected neural cultures at 24 and 72 hours post infection. Even though SARS-CoV-2 infection was scarce, IFNλ2/3 was induced in both the mixed neural culture and Ngn2 co-cultures, but not in NPC cultures. Increased secretion of IL-8 was observed in NPC cultures, mixed neural culture and Ngn2 co-cultures. H5N1 virus infection induced both type-III-IFN -IFNλ1 and -IFNλ2/3 in mixed neural cultures and Ngn2 co-cultures, but not among NPCs. Furthermore, increased levels of IP-10 were detected only in the H5N1 virus infected neural cultures. Similar to SARS-CoV2, H5N1 virus was also able to induce IL-8 in all neural cultures. Neither SARS-CoV-2, nor H5N1, virus infection induced type-I-interferon (IFNα/IFNβ) or type-II-IFN (IFNγ) or IL-1b, TNF-a, IL-12p70, GM-CSF or IL-10 (Supplement Figure 3).

## Discussion

SARS-CoV-2 replicated poorly in all three hiPSC-derived neural cultures used in our experiments, which contrasts largely to H5N1 virus, which replicated efficiently to high titers. Even though important entry factors for SARS-CoV-2 are expressed in all of the cultures used, SARS-CoV-2 infected a very small proportion of cells without evidence of subsequent spread or cellular apoptosis. However, SARS-CoV-2 infection did induce type-III-IFN and IL-8 production.

Evidence is accumulating that SARS-CoV-2 enters the CNS via the olfactory nerve^7–9^, a pathway that is also used by influenza A viruses to enter the CNS in many mammals including humans^10,37^. H5N1 virus spreads efficiently to the CNS via the olfactory nerve in experimentally inoculated ferrets and subsequently replicates very efficiently in the CNS^27,29,38^. Unlike H5N1 virus infection, SARS-CoV-2 is rarely detected in the CNS of fatal COVID-19 patients or experimentally inoculated animals^11–15^. In addition, only a handful of case reports of SARS-CoV-2 induced encephalitis have been reported^39,40^.. Altogether, these observations are consistent with our findings of poor SARS-CoV-2 replication in hiPSC-derived NPCs, neurons and astrocytes, and supports a pathophysiological model whereby SARS-CoV-2 invades the CNS, but does not replicate efficiently in CNS cell types. However, one caveat of our study is that other cells such as microglia, oligodendrocytes and vascular cells (pericytes, endothelial cells) are not present. Therefore, we cannot exclude that SARS-CoV-2 can infect and possibly replicate efficiently in other cells of the CNS or neuronal cell types such as cortical PV interneurons, midbrain or hindbrain cell types.

Despite the low proportion of SARS-CoV-2 infected cells and the fact that infection seemed to be abortive in the hiPSC derived neural cultures, we found evidence for cellular immune activation. In particular, SARS-CoV-2 infection of the neural cultures resulted in the induction of type-III-IFN and IFNλ2/3, but not type-I-IFN or type-II-IFN. This result is in accordance with earlier reports suggesting that SARS-CoV-2 triggers only very mild type-I and type-II-IFN responses, but does trigger a robust type-III-IFN response in cell culture, human airway epithelial cells, ferrets and SARS-CoV-2 infected individuals^41,42^. In addition, IL-8—a chemotactic factor that attracts leukocytes—was induced in all hiPSC-derived cultures. In lung tissue and peripheral venous blood serum of SARS-CoV-2 infected patients, elevated levels of IL-8 are associated with severe COVID-19^43–45^. Furthermore, IL-8 has been detected in the CSF of SARS-CoV-2 patients who developed encephalitis, which might be induced by the SARS-CoV-2 associated brain immune response, since SARS-CoV-2 RNA could not be detected in the CSF^46^. However, how exactly these cytokines contribute to the *in vivo* neuroinflammatory process, and if they are directly triggered by SARS-CoV-2 entry into the CNS needs further investigations

Highly pathogenic H5N1 virus replication has been reported *in vivo*^27,29,31,32,47^ and *in vitro* across several different types of human and mouse neural cell cultures^34–36^, including the human neuroblastoma line SK-N-SH^26^, suggesting this virus is neurotropic. This fits with our observation that H5N1 virus replicates productively and spreads throughout hiPSC-derived neural cultures, infecting NPCs as well as mature neurons and astrocytes. H5N1 virus infection also results in the upregulation of type-III-IFN, IFNλ1 and IFNλ2/3, as well as the antiviral cytokines IL-8 and IP-10. IP-10 has been detected in the CSF of influenza A virus infected patients and was found to be elevated in the brains of mice experimentally infected with H5N1 virus^48^. However, the mechanism by which H5N1 virus achieves abundant virus replication and robust induction of pro-neuroinflammatory cytokines remains poorly understood.

Altogether, our findings reveal that replication of SARS-CoV-2 in CNS cell types is very limited, which is in contrast to the efficient replication and spread of H5N1 virus. Although the mechanistic pathogenesis of SARS-CoV-2 associated CNS disease remains poorly understood, this study supports the hypothesis that SARS-CoV-2 entry into the CNS and direct infection of a small subset of neurons might trigger inflammation in the brain

## Material and Methods

### Cell Lines

VeroE6 (ATCC® CRL 1586TM) cells were maintained in Dulbecco’s modified Eagle’s medium (DMEM, Lonza, Breda, the Netherlands) supplemented with 10% fetal calf serum (FCS, Sigma-Aldrich, St. Louis, MO, USA), 10mM HEPES, 1.5 mg/ml sodium bicarbonate, 100 IU/ml penicillin (Lonza, Basel, Switzerland) and 100 μg/ml streptomycin (Lonza). Madin-Darby Canine Kidney (MDCK) cells were maintained in Eagle minimal essential medium (EMEM; Lonza) supplemented with 10% FCS, 100 IU/ml penicillin, 100 μg/ml streptomycin, 2 mM glutamine, 1.5 mg/ml sodium bicarbonate, 1 mM, 10 mM HEPES and 0.1 mM nonessential amino acids. All cell lines were grown at 37 °C in 5% CO_2_. The medium was refreshed every 3–4 days, and cells were passaged at >90% confluence with the use of PBS and trypsin-EDTA (0.05%). The cells were routinely checked for the presence of mycoplasma.

### Differentiation of iPSCs to NPCs and mature neural cultures

Human induced pluripotent stem cells (iPSCs) [WTC-11 Coriell #GM25256, obtained from the Gladstone Institute, San Francisco, USA] were differentiated to NPCs as previously described^2^ with slight modifications. After passage 3, NPC cultures were purified using fluorescence-activated cell sorting (FACS) as described previously^49^. Briefly, NPCs were detached from the culture plate and resuspended into a single cell solution. CD184+/CD44-/CD271-/CD24+ cells were collected using a FACSaria III (BD bioscience) and expanded in NPC medium (Table 2). NPCs were used for experiments between passage 3 and 7 after sorting or differentiated to neural networks. For differentiation towards mature neural cultures, NPCs were grown in neural differentiation medium (Table 2) for 6-8 weeks to achieve mature neural networks ^24^ and subsequently used for experiments, after week 4 only half of the medium was refreshed. Cultures were kept at 37 °C/5%CO2 throughout the entire differentiation process.

**Table 1.**
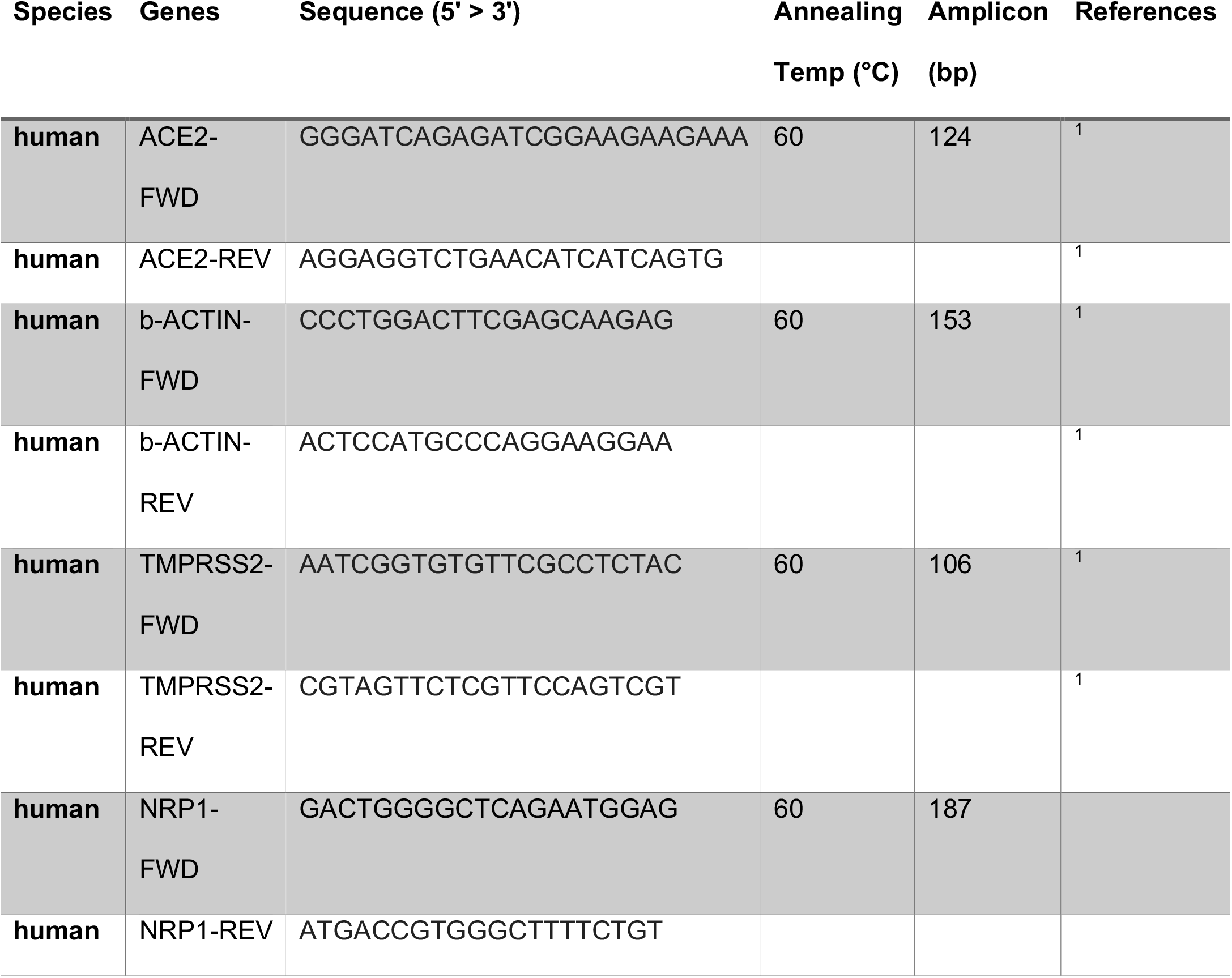

**Table 2.**
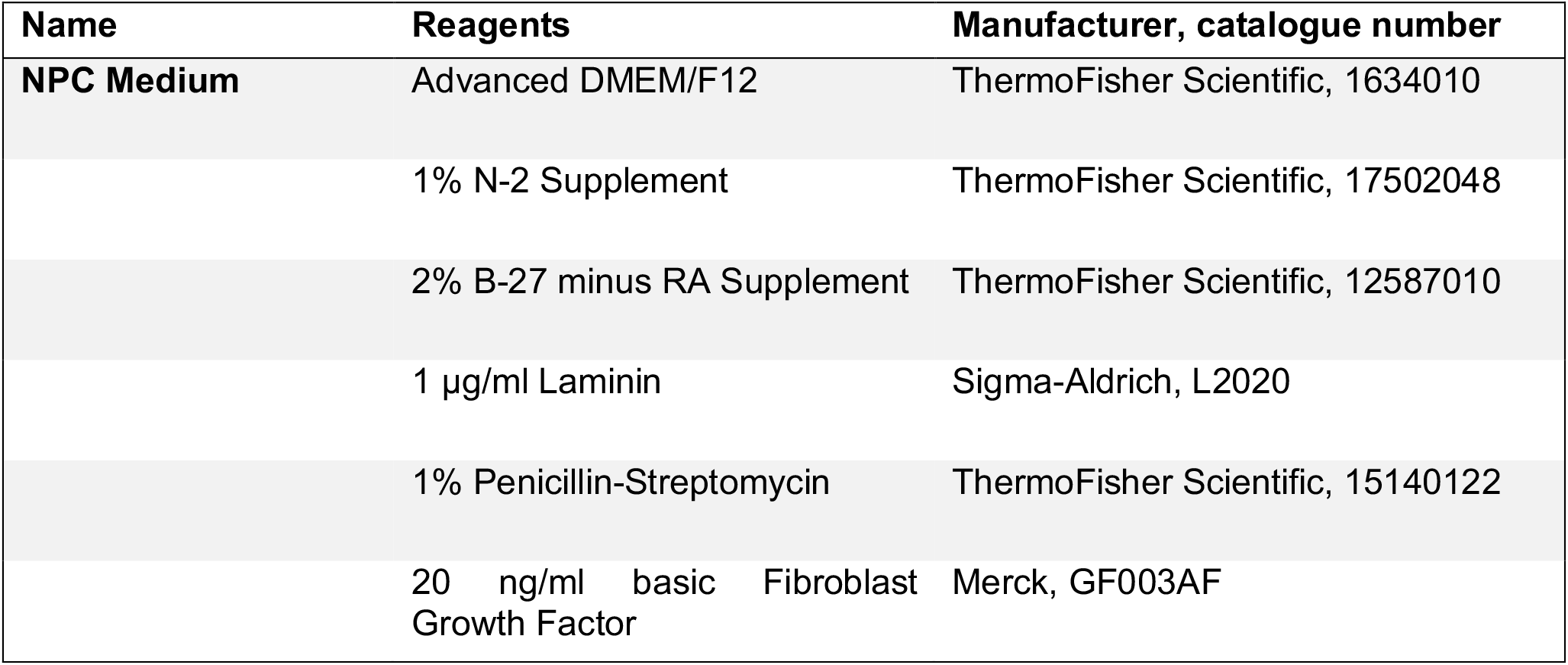
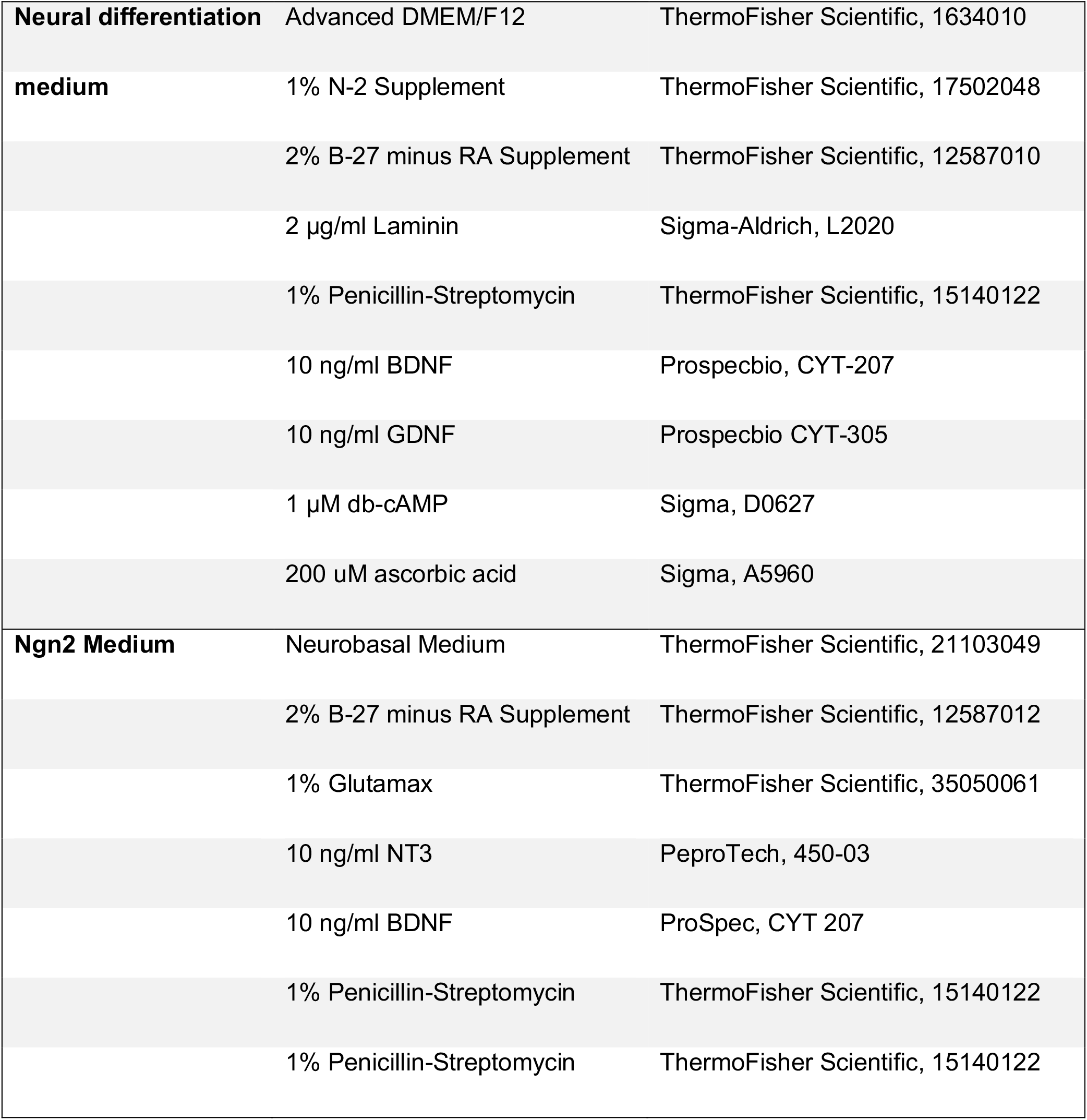
Overview of media and reagents used

**Table 3.** Antibodies

| **Antibody** | **Dilution** | **Manufacturer, catalogue number** |
| --- | --- | --- |
| SARS-CoV-2- Anti NP | 1:500 | Sino Biological, 40143-MM05 |
| SOX2 | 1:250 | Millipore, AB5603 |
| NEUN | 1:100 | Merck, ABN78 |
| GFAP | 1:200 | Millipore, AB5804 |
| MAP2 | 1:200 | Synaptic Systems, 188004 |
| H5N1- Anti NP | 1:1000 | EVL, EBS-I-047, Clone Hb65 |
| Caspase-3 | 1:100 | Cell Signaling Technologies, 9661S |

### Differentiation of iPSCs to Ngn2 co-cultures

iPSCs were directly differentiated into excitatory cortical layer 2/3 neurons by forcibly overexpressing the neuronal determinant Neurogenin 2 (Ngn2)^25,50^. To support neuronal maturation, hiPSC-derived astrocytes were added to the culture in a 1:1 ratio. At day 3, the medium was changed to Ngn2-medium (Table 2). Cytosine b-D-arabinofuranoside (Ara-C) (2 µM; Sigma, C1768) was added once to remove proliferating cells from the culture and ensure long-term recordings of the cultures. From day 6 onwards, half of the medium was refreshed three times per week. Cultures were kept at 37 °C/5%CO2 throughout the entire differentiation process.

### Viruses

The SARS-CoV-2 isolate (isolate BetaCoV/Munich/BavPat1/2020; European Virus Archive Global #026V-03883; kindly provided by Dr. C. Drosten) was previously described by *Lamers et al*.^51,52^ The zoonotic HPAI H5N1 virus (A/Indonesia/5/2005) was isolated from a human patient and the virus was propagated once in embryonated chicken eggs and twice in MDCK cells.

### Virus Titrations

The SARS-CoV-2 titers were determined by endpoint dilution on VeroE6 cells, calculated according to the method of Spearman & Kärber and expressed as 50% tissue culture infectious dose/ml (TCID_50_/ml). SARS-CoV-2 virus titers were determined by preparing 10-fold serial dilutions in triplicates of supernatants in Opti-MEM containing GlutaMAX. Dilutions supernatants were added to a monolayer of 40.000 VeroE6 cells/ well in a 96-well plateand incubate at 37°C. After 5 days the plates were examined for the presence of cytopathic effect (CPE). Virus titers of HPAI H5N1 were determined by endpoint dilution on MDCK as described previously^53^. In short, 10-fold serial dilutions of cell supernatant in triplicates were prepared in Influenza infection medium which consists of EMEM supplemented with 100 IU/ml penicillin, 100 μg/ml streptomycin, 2 mM glutamine, 1.5 mg/ml sodium bicarbonate, 10 mM HEPES, 1× (0.1 mM) nonessential amino acids, and 1 μg/μl TPCK-treated trypsin (Sigma-Aldrich). Prior to adding the virus dilutions to the MDCK cells, the cells were washed once with plain EMEM-medium to remove residual FCS. 100 µl of the diluted supernatants was used to inoculate 30.000 MDCK cells/well in a 96-well plate. After one hour, the inoculum was removed and 200µl fresh influenza infection medium was added. Four days after infection, supernatants of the infected MDCK cells were tested for agglutination. 25µl of the supernatant were mixed with 75µl 0.33% turkey red blood cells and incubated for one hour at 4°C. The titers of infectious virus were calculated according to the method of Spearman & Kärber and expressed as TCID_50_/ml. All experiment with infectious SARS-CoV-2 and H5N1 virus were performed in a Class II Biosafety Cabinet under BSL-3 conditions at the Erasmus Medical Center. The initial 1:10 dilution of cell supernatant resulted in a detection limit of 10^1.5^ TCID_50_/ml.

### Replication Kinetic

Before infection of neural progenitors, network neurons and NGN2 neurons, supernatant was removed and cells were infected with SARS-CoV-2 and H5N1 virus at the indicated multiplicity of infection (MOI). As control for active virus replication, VeroE6 and MDCK cells were infected with SARS-CoV-2 and H5N1 virus, respectively. Before virus infection, the VeroE6 cells were washed with SARS-CoV-2 infection medium (DMEM supplemented with 2% FBS and 100 IU/ml penicillin, 100 μg/ml streptomycin) and MDCK cells were washed with influenza infection medium. After 1 hours of incubation at 37°C, the inoculum was removed and replaced with fresh medium and old medium in a 1:1 ratio. After removing the inoculum from VeroE6 and MDCK cells, SARS-CoV-2 infection medium and influenza infection medium, respectively, were added to the cells. At the indicated timepoints an aliquot of the supernatant was collected for subsequent virus titration. All experiments were performed in biological triplicates and either in technical duplicates or triplicates.

### PCR Validation of ACE2, TMPRSS2 and NRP1 expression

RNA was isolated from the neural cultures and VeroE6 cells using the High Pure RNA Isolation Kit (Roche). The concentration of RNA was determined using Nanodrop. 2.5µg of RNA was reverse transcribed into cDNA using the SuperScript III reverse transcriptase (Invitrogen) according to the manufacturers’ protocol. cDNA of bronchus-bronchiol organoids were kindly provided by Anna Z. Mykytyn. Presence of ACE2, TMPRSS2, NRP1 and ACT was evaluated by amplified these genes with gene specific primers (Table 1) by PCR. Gene products were visualized on a 2% agarose gel which was stained with SYBR Safe. PCR products of the genes were sequenced to validate that the right product was amplified.

### Multiplexed bead-assay for Cytokine Profiling

Cytokines were measured using the LEGENDplex™ Human Anti-Virus Response Panel (BioLegend). The kit was used according to manufacturer’s manual with an additional fixing step. After adding the SA-PE and performing the washing steps, the supernatant and the beads were fixed with formalin for 15 minutes at room temperature and washed twice with the provided wash buffer. This ensures that all pathogens are not infectious.

### Immunofluorescent labeling

Cells were fixed using 4% formalin in PBS and labeled using immunocytochemistry. Primary antibody incubation was performed overnight at 4°C. Secondary antibody incubation was performed for 2 hours at room temperature. Both primary and secondary antibody incubation was performed in staining buffer [0.05 M Tris, 0.9% NaCl, 0.25% gelatin, and 0.5% Triton X-100 (Sigma, T8787) in PBS (pH 7.4)]. Primary antibodies and their dilutions can be found in Table 2. Secondary antibodies conjugated to Alexa-488, Alexa-647 or Cy3 were used at a dilution of 1:400 (Jackson ImmunoResearch). Nuclei were visualised using DAPI (ThermoFisher Scientific, D1306). Samples were embedded in Mowiol 4-88 (Sigma-Aldrich, 81381) and imaged using a Zeiss LSM 800 confocal microscope (Oberkochen, Germany).

## Acknowledgments

We thank Anna Z. Mykytyn and Mart M. Lamers for sharing reagents, cDNA of the airway organoids, SARS-CoV-2 virus stocks and for technical advice and scientific discussions. This work was funded by a fellowship to D.V.R. from the Netherlands Organization for Scientific Research (VIDI contract 91718308) and a EUR fellowship. This work was also supported by the Netherlands Organ-on-Chip Initiative, an NWO Gravitation project (024.003.001) funded by the Ministry of Education, Culture and Science of the government of the Netherlands (S.K, F.D.V., B.L.) and by an Erasmus MC Human Disease Model Award to F.D.V.

The authors declare no conflict of interest.

## Notes

### Competing Interest Statement

The authors have declared no competing interest.

## References

1. Varatharaj, A. et al. Neurological and neuropsychiatric complications of COVID-19 in 153 patients: a UK-wide surveillance study. The Lancet Psychiatry 7, 875–882 (2020).

2. Bryche, B. et al. Massive transient damage of the olfactory epithelium associated with infection of sustentacular cells by SARS-CoV-2 in golden Syrian hamsters. Brain. Behav. Immun. 89, 579–586 (2020).

3. Harapan, B. N. & Yoo, H. J. Neurological symptoms, manifestations, and complications associated with severe acute respiratory syndrome coronavirus 2 (SARS-CoV-2) and coronavirus disease 19 (COVID-19). J. Neurol. 2, (2021).

4. Guadarrama-Ortiz, P. et al. Neurological Aspects of SARS-CoV-2 Infection: Mechanisms and Manifestations. Front. Neurol. 11, 1–14 (2020).

5. Solomon, I. H. et al. Neuropathological Features of Covid-19. N. Engl. J. Med. (2020) doi:10.1056/nejmc2019373.

6. Pennisi, M. et al. Sars-cov-2 and the nervous system: From clinical features to molecular mechanisms. Int. J. Mol. Sci. 21, 1–21 (2020).

7. Zou, L. et al. SARS-CoV-2 Viral Load in Upper Respiratory Specimens of Infected Patients. N. Engl. J. Med. 382, 1175–1177 (2020).

8. Sia, S. F. et al. Pathogenesis and transmission of SARS-CoV-2 in golden hamsters. Nature 583, 834–838 (2020).

9. Meinhardt, J. et al. Olfactory transmucosal SARS-CoV-2 invasion as a port of central nervous system entry in individuals with COVID-19. Nat. Neurosci. 24, 168–175 (2021).

10. Riel, D. Van, Verdijk, R. & Kuiken, T. The olfactory nerve: A shortcut for influenza and other viral diseases into the central nervous system. J. Pathol. 235, 277–287 (2015).

11. Matschke, J. et al. Neuropathology of patients with COVID-19 in Germany: a postmortem case series. Lancet Neurol. 19, 919–929 (2020).

12. Schurink, B. et al. Viral presence and immunopathology in patients with lethal COVID-19: a prospective autopsy cohort study. The Lancet Microbe 1, e290–e299 (2020).

13. Edén, A. et al. CSF Biomarkers in Patients With COVID-19 and Neurologic Symptoms: A Case Series. Neurology 96, e294–e300 (2021).

14. Neumann, B. et al. Cerebrospinal fluid findings in COVID-19 patients with neurological symptoms. J. Neurol. Sci. 418, (2020).

15. Destras, G. et al. Systematic SARS-CoV-2 screening in cerebrospinal fluid during the COVID-19 pandemic. The Lancet Microbe 1, e149 (2020).

16. Dobrindt, K. et al. Common genetic variation in humans impacts in vitro susceptibility to SARS-CoV-2 infection. Stem Cell Reports 16, 1–14 (2021).

17. Pellegrini, L. et al. SARS-CoV-2 Infects the Brain Choroid Plexus and Disrupts the Blood-CSF Barrier in Human Brain Organoids. Cell Stem Cell 27, 951-961.e5 (2020).

18. Zhang, B. Z. et al. SARS-CoV-2 infects human neural progenitor cells and brain organoids. Cell Res. 30, 928–931 (2020).

19. Bullen, C. K. et al. Infectability of human BrainSphere neurons suggests neurotropism of SARS-CoV-2. ALTEX 37, 665–671 (2020).

20. Wang, C. et al. ApoE-Isoform-Dependent SARS-CoV-2 Neurotropism and Cellular Response. Cell Stem Cell 28, 331-342.e5 (2021).

21. Song, E. et al. Neuroinvasion of SARS-CoV-2 in human and mouse brain. J. Exp. Med. 218, (2021).

22. Ramani, A. et al. SARS -CoV-2 targets neurons of 3D human brain organoids. EMBO J. 39, 1–14 (2020).

23. Jacob, F. et al. Human Pluripotent Stem Cell-Derived Neural Cells and Brain Organoids Reveal SARS-CoV-2 Neurotropism Predominates in Choroid Plexus Epithelium. Cell Stem Cell 27, 937-950.e9 (2020).

24. Gunhanlar, N. et al. A simplified protocol for differentiation of electrophysiologically mature neuronal networks from human induced pluripotent stem cells. Mol. Psychiatry 23, 1336–1344 (2018).

25. Zhang, Y. et al. Rapid single-step induction of functional neurons from human pluripotent stem cells. Neuron 78, 785–798 (2013).

26. Frega, M. et al. Rapid Neuronal Differentiation of Induced Pluripotent Stem Cells for Measuring Network Activity on Micro-electrode Arrays. J. Vis. Exp. (2017) doi:10.3791/54900.

27. Schrauwen, E. J. A. et al. The Multibasic Cleavage Site in H5N1 Virus Is Critical for Systemic Spread along the Olfactory and Hematogenous Routes in Ferrets. J. Virol. 86, 3975–3984 (2012).

28. Shinya, K. et al. Subclinical Brain Injury Caused by H5N1 Influenza Virus Infection. J. Virol. 85, 5202–5207 (2011).

29. Bodewes, R. et al. Pathogenesis of influenza A/H5N1 virus infection in ferrets differs between intranasal and intratracheal routes of inoculation. Am. J. Pathol. 179, 30–36 (2011).

30. Park, C. H. et al. The invasion routes of neurovirulent A/Hong Kong/483/97 (H5N1) influenza virus into the central nervous system after respiratory infection in mice. Arch. Virol. 147, 1425–1436 (2002).

31. Jang, H. et al. Highly pathogenic H5N1 influenza virus can enter the central nervous system and induce neuroinflammation and neurodegeneration. Proc. Natl. Acad. Sci. U. S. A. 106, 14063–14068 (2009).

32. Shinya, K. et al. Systemic Dissemination of H5N1 Influenza A Viruses in Ferrets and Hamsters after Direct Intragastric Inoculation. J. Virol. 85, 4673–4678 (2011).

33. Siegers, J. Y. et al. Viral Factors Important for Efficient Replication of Influenza A Viruses in Cells of the Central Nervous System. J. Virol. 93, 1–12 (2019).

34. Ng, Y. P. et al. Avian influenza H5N1 virus induces cytopathy and proinflammatory cytokine responses in human astrocytic and neuronal cell lines. Neuroscience 168, 613–623 (2010).

35. Pringproa, K. et al. Tropism and induction of cytokines in human embryonic-stem cells-derived neural progenitors upon inoculation with highly-pathogenic avian H5N1 influenza virus. PLoS One 10, 1–14 (2015).

36. Lin, X. et al. Insights into human astrocyte response to H5N1 infection by microarray analysis. Viruses 7, 2618–2640 (2015).

37. Van Riel, D. et al. Evidence for influenza virus CNS invasion along the olfactory route in an immunocompromised infant. J. Infect. Dis. 210, 419–423 (2014).

38. Shinya, K. et al. Avian influenza virus intranasally inoculated infects the central nervous system of mice through the general visceral afferent nerve. Arch. Virol. 145, 187–195 (2000).

39. Casez, O. et al. Teaching NeuroImages: SARS-CoV-2-Related Encephalitis: MRI Pattern of Olfactory Tract Involvement. Neurology 96, e645–e646 (2021).

40. Vandervorst, F. et al. Encephalitis associated with the SARS-CoV-2 virus: A case report. Interdiscip. Neurosurg. Adv. Tech. Case Manag. 22, 0–2 (2020).

41. Vanderheiden, A. et al. Type I and Type III Interferons Restrict SARS-CoV-2 Infection of Human Airway Epithelial Cultures. J. Virol. 94, 1–16 (2020).

42. Blanco-Melo, D. et al. Imbalanced Host Response to SARS-CoV-2 Drives Development of COVID-19. Cell 181, 1036-1045.e9 (2020).

43. Del Valle, D. M. et al. An inflammatory cytokine signature predicts COVID-19 severity and survival. Nat. Med. 26, 1636–1643 (2020).

44. van Riel, D. et al. Temporal kinetics of RNAemia and associated systemic cytokines in hospitalized COVID-19 patients. bioRxiv (2020) doi:10.1101/2020.12.17.423376.

45. Chen, L. et al. Scoring cytokine storm by the levels of MCP-3 and IL-8 accurately distinguished COVID-19 patients with high mortality. Signal Transduct. Target. Ther. 5, 8–10 (2020).

46. Benameur, K. et al. Encephalopathy and encephalitis associated with cerebrospinal fluid cytokine alterations and coronavirus disease, atlanta, georgia, usa, 2020. Emerg. Infect. Dis. 26, 2016–2021 (2020).

47. Park, C. H. et al. Persistence of viral RNA segments in the central nervous system of mice after recovery from acute influenza A virus infection. Vet. Microbiol. 97, 259–268 (2003).

48. Jang, H. et al. Inflammatory effects of highly pathogenic H5N1 influenza virus infection in the CNS of mice. J. Neurosci. 32, 1545–1559 (2012).

49. Yuan, S. H. et al. Cell-surface marker signatures for the Isolation of neural stem cells, glia and neurons derived from human pluripotent stem cells. PLoS One (2011) doi:10.1371/journal.pone.0017540.

50. Frega, M. et al. Rapid neuronal differentiation of induced pluripotent stem cells for measuring network activity on micro-electrode arrays. J. Vis. Exp. (2017) doi:10.3791/54900.

51. Lamers, M. M. et al. SARS-CoV-2 productively infects human gut enterocytes. Science (80-.). 3, 50–54 (2020).

52. Lamers, M. M. et al. Human airway cells prevent SARS-CoV-2 multibasic cleavage site cell culture adaptation 1 2. bioRxiv (2021).

53. Rimmelzwaan, G. F., Baars, M., Claas, E. C. J. & Osterhaus, A. D. M. E. Comparison of RNA hybridization, hemagglutination assay, titration of infectious virus and immunofluorescence as methods for monitoring influenza virus replication in vitro. J. Virol. Methods (1998) doi:10.1016/S0166-0934(98)00071-8.

