## Supplement for "Replication kinetic, cell tropism and associated immune responses in SARS-CoV-2 and H5N1 virus infected human iPSC derived neural models"

### Supplement Figures

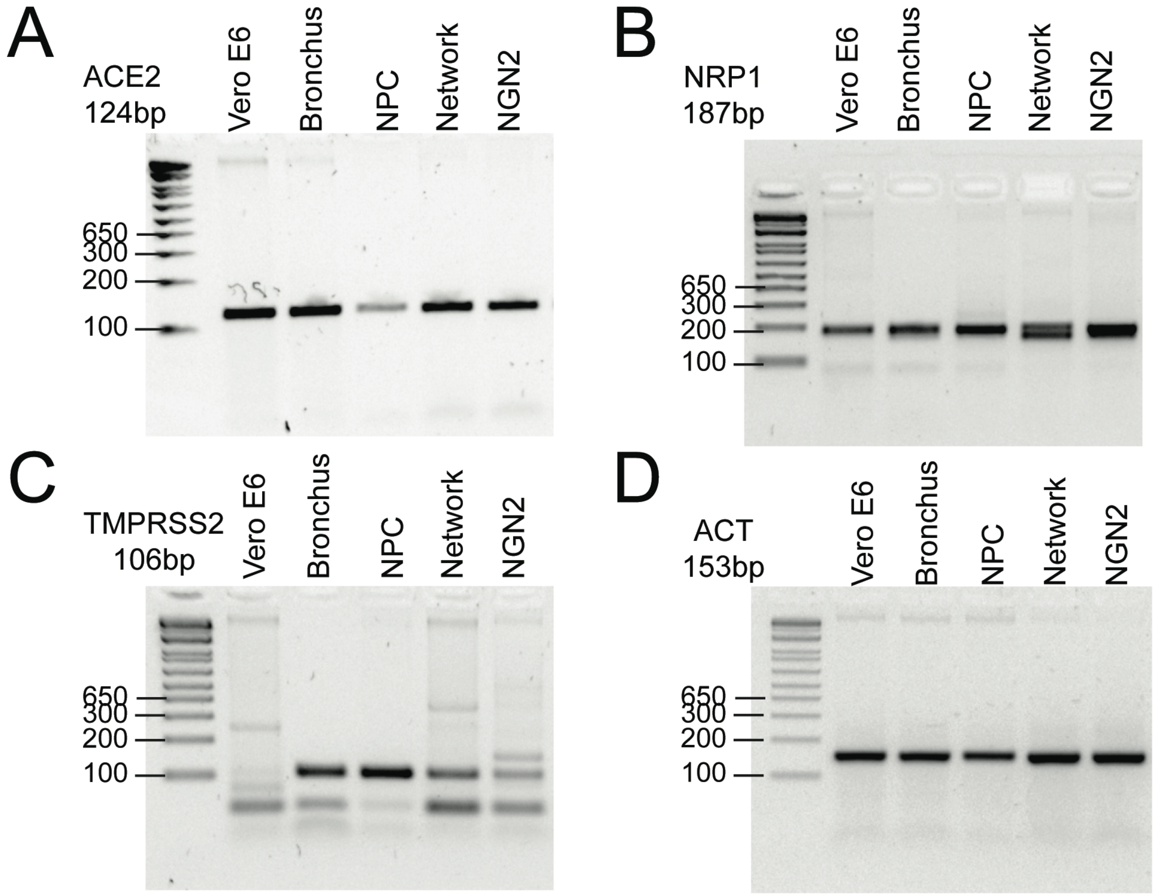


**Supplement Figure 1. Expression of important entry factors for SARS-CoV-2, original gel pictures.** Expression of (A) ACE2 with amplicon size 124bp (B) NRP1 with amplicon size 187 bp (C) TMPRSS2 with amplicon size 106bp (D) ACT with amplicon size 153bp from one out of two independent experiment are shown. The important bands with the correct sizes of the amplicon were cropped and are represented in Figure 1G.

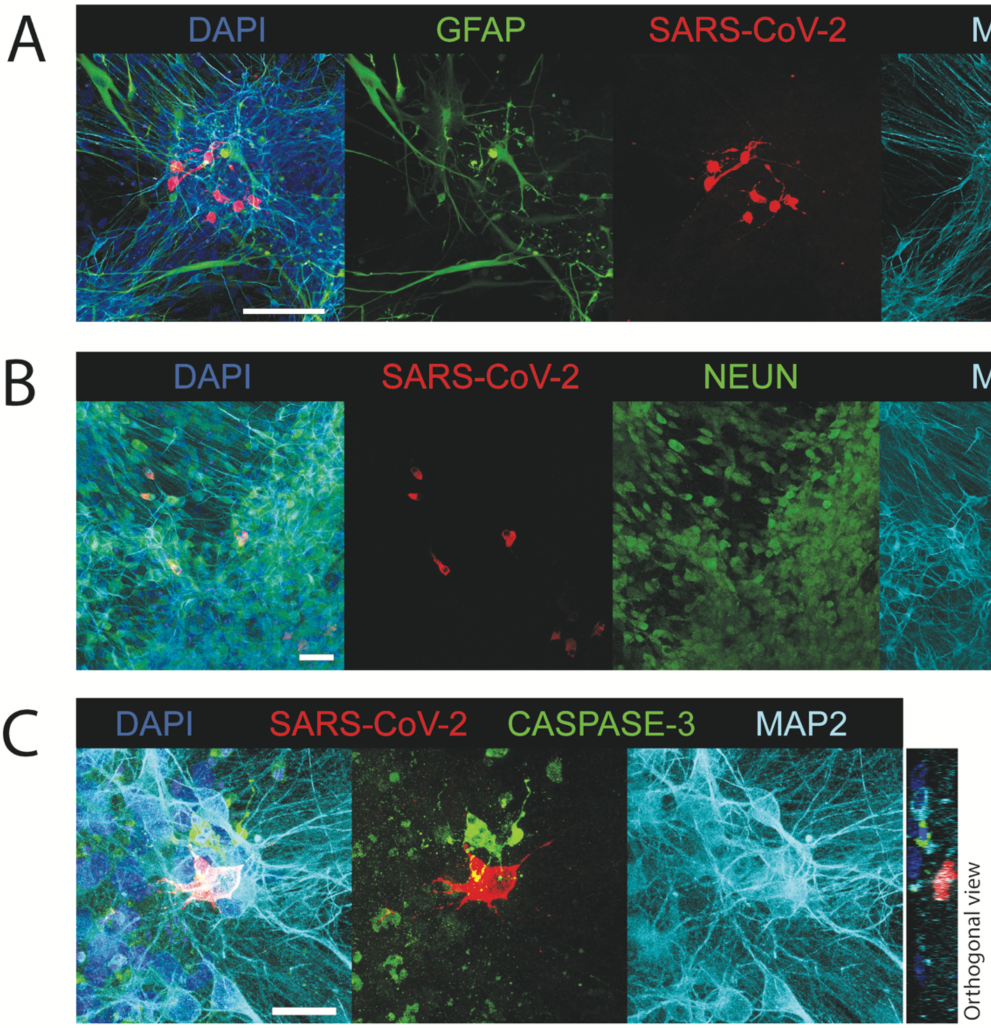


**Supplement Figure 2. SARS-CoV2 infects NEUN$^+$ neurons and does not induce caspase-3 expression.** (A) One focus of multiple infected cells was identified (scale bar = 100 µm). (B) The neurons in the Ngn2 cultures consist of a homogeneous population of mature NEUN$^+$ neurons (scale bar = 50 µm). (C) High magnification image of a SARS-CoV-2 infected cells in which no caspase-3 is visible (scale bar = 20 µm).

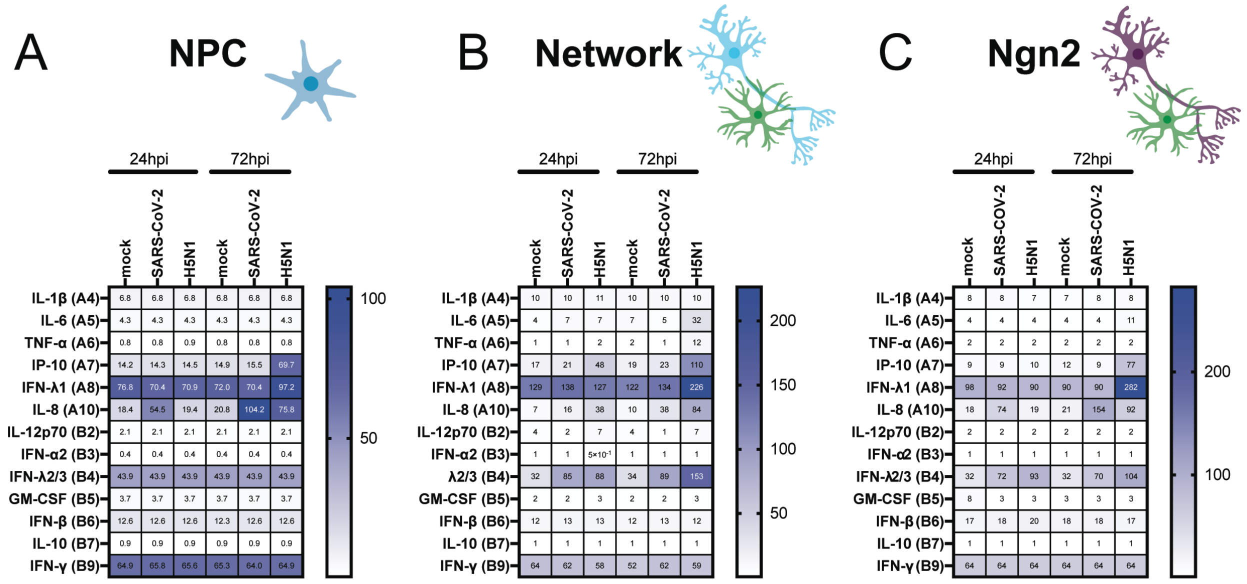


**Supplement Figure 3. SARS-CoV-2 infection as well as H5N1 virus infection results in modest immune responses.** The concentration of the cytokines which were measures in the supernatant of SARS-CoV-2 or H5N1 virus infected (A) NPCs, (B) mixed neural cultures and (C) Ngn2 co-cultures are displayed. The values represent the means of three independent experiments.
